# *De Novo* Assembly of the Northern Cardinal (*Cardinalis cardinalis*) Genome Reveals Candidate Regulatory Regions for Sexually Dichromatic Red Plumage Coloration

**DOI:** 10.1101/2020.05.12.092080

**Authors:** Simon Yung Wa Sin, Lily Lu, Scott V. Edwards

## Abstract

Northern cardinals (*Cardinalis cardinalis*) are common, mid-sized passerines widely distributed in North America. As an iconic species with strong sexual dichromatism, it has been the focus of extensive ecological and evolutionary research, yet genomic studies investigating the evolution of genotype–phenotype association of plumage coloration and dichromatism are lacking. Here we present a new, highly contiguous assembly for *C. cardinalis*. We generated a 1.1 Gb assembly comprised of 4,762 scaffolds, with a scaffold N50 of 3.6 Mb, a contig N50 of 114.4 kb and a longest scaffold of 19.7 Mb. We identified 93.5% complete and single-copy orthologs from an Aves dataset using BUSCO, demonstrating high completeness of the genome assembly. We annotated the genomic region comprising the CYP2J19 gene, which plays a pivotal role in the red coloration in birds. Comparative analyses demonstrated non-exonic regions unique to the CYP2J19 gene in passerines and a long insertion upstream of the gene in *C. cardinalis*. Transcription factor binding motifs discovered in the unique insertion region in *C. cardinalis* suggest potential androgen-regulated mechanisms underlying sexual dichromatism. Pairwise Sequential Markovian Coalescent (PSMC) analysis of the genome reveals fluctuations in historic effective population size between 100,000–250,000 in the last 2 millions years, with declines concordant with the beginning of the Pleistocene epoch and Last Glacial Period. This draft genome of *C. cardinalis* provides an important resource for future studies of ecological, evolutionary, and functional genomics in cardinals and other birds.

## Introduction

The northern cardinal (*Cardinalis cardinalis*) is a mid-sized (∼42-48 g) passerine broadly distributed in eastern and central North America, with a range encompassing northern Central America to southeastern Canada (Smith et al. 2011). It has high genetic and phenotypic diversity and is currently divided into 18 subspecies (Paynter 1970; Smith et al. 2011). Cardinals have been studied extensively on research areas such as song and communication (e.g. Anderson and Conner 1985; Yamaguchi 1998; Jawor and MacDougall-Shackleton 2008), sexual selection (e.g. Jawor et al. 2003), physiology (e.g. DeVries and Jawor 2013; Wright and Fokidis 2016), phylogeography (e.g. Smith et al. 2011), and plumage coloration (e.g. Linville and Breitwisch 1997; Linville et al. 1998; McGraw et al. 2003). *C. cardinalis* is an iconic species in Cardinalidae and has strong sexual dichromatism, with adult males possessing bright red plumage and adult females tan (Fig. 1).

**Figure 1.**
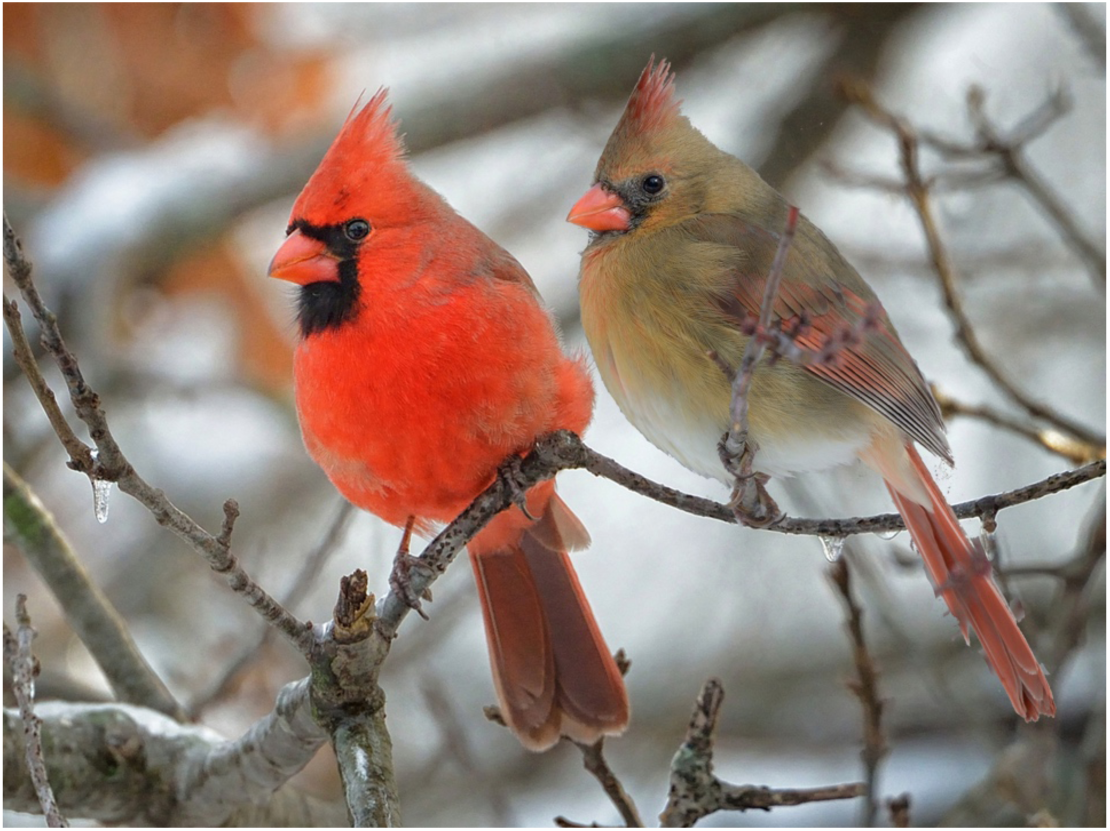
A pair of northern cardinals, a common, sexually dichromatic passerine bird. The adult male (left) has bright red plumage whereas the adult female (right) is primarily tan in color. Photo © Clarence Stewart.

The red plumage coloration of many bird species plays important roles in social and sexual signaling. With the advance of genomic technologies in the last several years, the genetic basis and evolution of plumage coloration in birds is under intense investigation (Lopes et al. 2016; Mundy et al. 2016; Twyman et al. 2016; Toomey et al. 2018; Twyman et al. 2018a; Twyman et al. 2018b; Funk and Taylor 2019; Kim et al. 2019; Gazda et al. 2020). The red color of feathers is generated by the deposition of ingested carotenoids, modified endogenously. A ketolase is involved in an oxidative reaction to convert dietary yellow carotenoids into red ketocarotenoids (Friedman et al. 2014). Recently the gene encoding the carotenoid ketolase has been shown to be a cytochrome P450 enzyme, CYP2J19 (Lopes et al. 2016; Mundy et al. 2016). The identification of CYP2J19 as the gene responsible for red coloration in hybrid canary (*Serinus canaria*) plumage (Lopes et al. 2016) and zebra finch (*Taeniopygia guttata*) bill and legs (Mundy et al. 2016) suggests its general role in red pigmentation of multiple tissues across birds. *C. cardinalis* is an excellent candidate to further our understanding of the regulation and development of red plumage coloration in birds. In particular, little attention has been paid to noncoding, regulatory signatures in the region surrounding CYP2J19, and genomic resources for cardinals would greatly facilitate such work.

Despite extensive ecological and evolutionary research on *C. cardinalis*, we lack a highly contiguous genome of this species and other species in the family Cardinalidae. A genome assembly of *C. cardinalis* will facilitate studies of genotype–phenotype association of sexually dichromatic plumage color in this species, and facilitate comparative genomic analysis in birds generally. Here we generate a new genome assembly from a male collected in Texas, USA, with a voucher specimen in the ornithology collection of the Museum of Comparative Zoology. We used the AllPaths-LG (Gnerre et al. 2011) method to perform *de novo* genome assembly. Given the lack of genomic resources currently available for the Cardinalidae, this *C. cardinalis* draft genome will facilitate future studies on genomic studies of cardinals and other passerines.

## Methods & Materials

### Sample collection and DNA extraction

We collected a male *C. cardinalis* on the DNA Works Ranch, Afton, Dickens County, Texas, United States (33.76286°, -100.8168°) on 19 January 2016 (MCZ Ornithology cat. no.: 364215). We collected muscle, heart, and liver and snap frozen the tissues in liquid nitrogen immediately in the field. Upon returning from the field, tissues were stored in the cryopreservation facility of the Museum of Comparative Zoology, Harvard University until DNA extraction. We isolated genomic DNA using the DNeasy Blood and Tissue Kit (Qiagen, Hilden, Germany) following the manufacturer’s protocol. We confirmed the sex of the individual using published PCR primers targeting CHD1 genes with different sizes of intron on the W and Z chromosomes (2550F & 2718R; Fridolfsson and Ellegren 1999) and measured DNA concentration with a Qubit dsDNA HS Assay Kit (Invitrogen, Carlsbad, USA).

### Library preparation and sequencing

We performed whole-genome library preparation and sequencing following Grayson et al. (2017). In brief, a DNA fragment library of 220 bp insert size was prepared using the PrepX ILM 32i DNA Library Kit (Takara), and mate-pair libraries of 3 kb insert size were prepared using the Nextera Mate Pair Sample Preparation Kit (cat. No. FC-132-1001, Illumina). We then assessed library quality using the HS DNA Kit (Agilent) and quantified the libraries with qPCR prior to sequencing (KAPA library quantification kit). We sequenced the libraries on an Illumina HiSeq instrument (High Output 250 kit, PE 125 bp reads) at the Bauer Core facility at Harvard University.

### De novo genome assembly and assessment

We assessed the quality of the sequencing data using FastQC and removed adapters using Trimmomatic (Bolger et al. 2014) and assembled the genome using AllPaths-LG v52488 (Gnerre et al. 2011), which allowed us to estimate the genome size from k-mer frequencies and assess the contiguity of the *de novo* genome. We estimated the completeness of the assembled genome with BUSCO v2.0 (Simão et al. 2015) and used the aves_odb9 dataset to search for 4915 universal single-copy orthologs in birds.

### Analysis of the CYP2J19 genomic region

We annotated the genomic region that comprises the CYP2J19 gene using blastn with the CYP2J19, CYP2J40, HOOK1 and NFIA genes from *S. canaria* and *T. guttata* as queries. Conserved non-exonic elements (CNEEs) were obtained from a published UCSC Genome Browser track hub containing a progressiveCactus alignment of 42 bird and reptile species and CNEE annotations (viewed in the UCSC genome browser at https://ifx.rc.fas.harvard.edu/pub/ratiteHub8/hub.txt; Sackton et al. 2019). We identified CYP2J19 genes from 30 bird species (Twyman et al. 2018a) in the NCBI by blasting their genome assemblies. The alignment was compiled using MUSCLE (Edgar 2004) and viewed with Geneious (Kearse et al. 2012). We also aligned the genomic regions up- and downstream of CYP2J19 in 10 passerines to identify any potential conserved regulatory regions present in species with red carotenoid coloration.

We used the MEME Suite (Bailey et al. 2009; Bailey et al. 2015) to discover potential regulatory DNA motifs in the region upstream of the CYP2J19 gene that we found to be unique to *C. cardinalis* and to predict transcription factors (TFs) binding to those motifs. We used MEME (Bailey and Elkan 1994) to identify the top five most statistically significant motifs and their corresponding positions in the region. We used Tomtom (Gupta et al. 2007) to compare the identified motifs against databases (e.g. JASPAR) of known motifs and identify potential TFs specific to those matched motifs. To search for potential biological roles of these motifs, we used GOMo (Buske et al. 2010) to scan all promoters in *Gallus gallus* using the identified motifs to determine if any motifs are significantly associated with genes linked to any Genome Ontology (GO) terms.

### Inference of demographic history

To investigate historical demographic changes in the northern cardinal we used the Pairwise Sequential Markovian Coalescent (PSMC) model (Li and Durbin 2011) based on the diploid whole-genome sequence to reconstruct the population history. We generated consensus sequences for all autosomes using SAMtools’ (v1.5; Li et al. 2009) *mpileup* command and the *vcf2fq* command from vcfutils.pl. We applied filters for base quality and mapping quality below 30. The settings for the PSMC atomic time intervals were “4+30*2+4+6+10”. 100 bootstraps were used to compute the variance in estimates of *N*_*e*_. To convert inferred population sizes and times to numbers of individuals and years, respectively, we used the estimate of mutation rate of 3.44e-09 per site per generation from the medium ground finch (*Geospiza fortis*, Thraupidae, a closely related passerine clade) (Nadachowska-Brzyska et al. 2015). We estimated the generation time of the northern cardinal as age of sexual maturity multiplied by a factor of two (Nadachowska-Brzyska et al. 2015), yielding a generation time of 2 years.

### Data availability (Data will be available later prior to publication)

Data from all sequencing runs and final genome assembly are available from NCBI (BioProject number will be provided later). Short-fragment and mate-pair libraries are also available from the NCBI SRA (number to be provided later).

## Results and Discussion

### Genome assembly and evaluation

We generated 703,754,970 total reads from two different sequencing libraries, including 335,713,090 reads from the fragment library and 368,041,880 reads from the mate-pair library. The genome size estimated by AllPaths-LG from k-mers was 1.1 Gb (Table 1). The sequence coverage estimated was ∼59x. The assembly consisted of 32,783 contigs and 4,762 scaffolds. The largest scaffold was 19.7 Mb. The contig N50 was 114.4 kb and the scaffold N50 was 3.6 Mb (Table 1). The number of contigs per Mb was 31.4 and the number of scaffolds per Mb was 4.6. The average GC content of the assembly was 42.1%. BUSCO scores (Simão et al. 2015) suggest high completeness of the genome, with 93.5% of single-copy orthologs for birds identified (Table 2). The genome contiguity and completeness is better than most genomes presented in Jarvis et al. (2014) and Grayson et al. (2017).

**Table 1.** *De novo* assembly metrics for northern cardinal genome

| Metric | Value |
| --- | --- |
| Estimated genome size | 1.10 Gb |
| %GC content | 42.1 |
| Total depth of coverage | 59x |
| Total contig length (bp) | 1,019,501,986 |
| Total scaffold length (bp, gapped) | 1,044,184,327 |
| Number of contigs | 32,783 |
| Contig N50 | 114.4 kb |
| Number of scaffolds | 4,762 |
| Scaffold N50 (with gaps) | 3.6 Mb |
| Largest scaffold | 19.7 Mb |

**Table 2.** Output from BUSCO analyses to assess genome completeness by searching for single-copy orthologs from aves dataset

|  | Aves | % |
| --- | --- | --- |
| Complete BUSCOs | 4642 | 94.4 |
| Complete and single-copy BUSCOs | 4596 | 93.5 |
| Complete and duplicated BUSCOs | 46 | 0.9 |
| Fragmented BUSCOs | 167 | 3.4 |
| Missing BUSCOs | 106 | 2.2 |
| Total BUSCO groups searched | 4915 |  |

### Candidate non-coding regulatory regions of CYP2J19

The CYP2J19 gene was identified on scaffold 100 of the *C. cardinalis* genome, flanked by CYP2J40 and NFIA genes (Fig. 2), an arrangement consistent with other species (e.g. Lopes et al. 2016; Mundy et al. 2016). The length of the CYP2J19 gene in *C. cardinalis* is 8925 bp, comprising 9 exons. We identified 26 CNEEs in the CYP2J19 gene, 24 CNEEs within ∼6 kb upstream and 1 CNEE within ∼6 kb downstream of the gene. Of those 51 CNEEs, 10 CNEEs in CYP2J19 and 7 CNEEs upstream of the gene are at least 50 bp in length.

**Figure 2.**
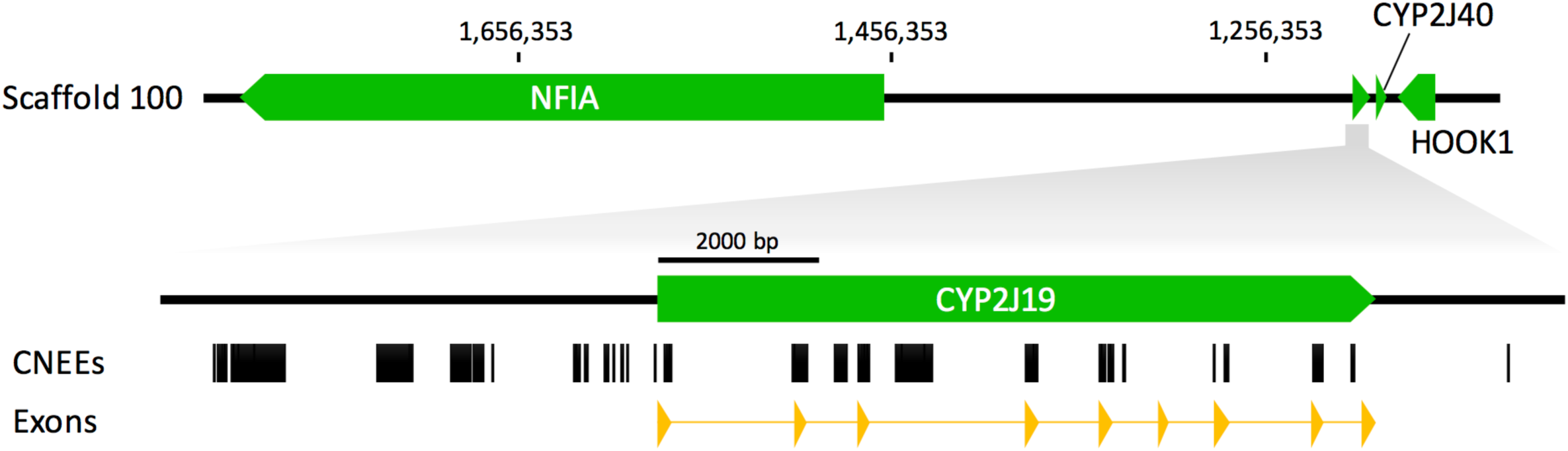
Annotation of the CYP2J19 region in scaffold 100 of the northern cardinal genome. The CYP2J19 gene is flanked by the NFIA and CYP2J40 genes as in other passerines. Conserved non-exonic elements (CNEEs) are shown in black boxes. Exons of the CYP2J19 gene are in yellow triangles.

Three intronic insertions, as well as one non-coding region upstream and one downstream of the gene were found to be present only in passerine species (Fig. 3). In addition, we identified unique insertions of large genomic regions in the two passerine species in our alignment with red carotenoid coloration, i.e. *C. cardinalis* and *T. guttata*. A unique 5920 bp insertion is present 339 bp upstream of the CYP2J19 gene in *C. cardinalis* (Fig. 4a). There is a 11322 bp insertion downstream of CYP2J19 in *T. guttata* comprising the CYP2J19B gene (Fig. 4b; Mundy et al. 2016) that is not present in *C. cardinalis*. The most similar sequence to the *C. cardinalis* upstream insertion identified by blasting (∼84% identity, 25% query cover) is a sequence annotated as “RNA-directed DNA polymerase from mobile element jockey-like” in the American crow (*Corvus brachyrhynchos*) genome assembly (Zhang et al. 2014), suggesting that the insertion may contain a non-long terminal repeat (non-LTR) retrotransposon (Ivanov et al. 1991). An endogenous retroviral insertion located upstream of the aromatase gene is proposed to be the mechanism of gene activation that lead to the henny feathers phenotype in chickens (Matsumine et al. 1991). The possibility that the upstream retrovirus may promote CYP2J19 gene activation is worth more investigation.

**Figure 3.**
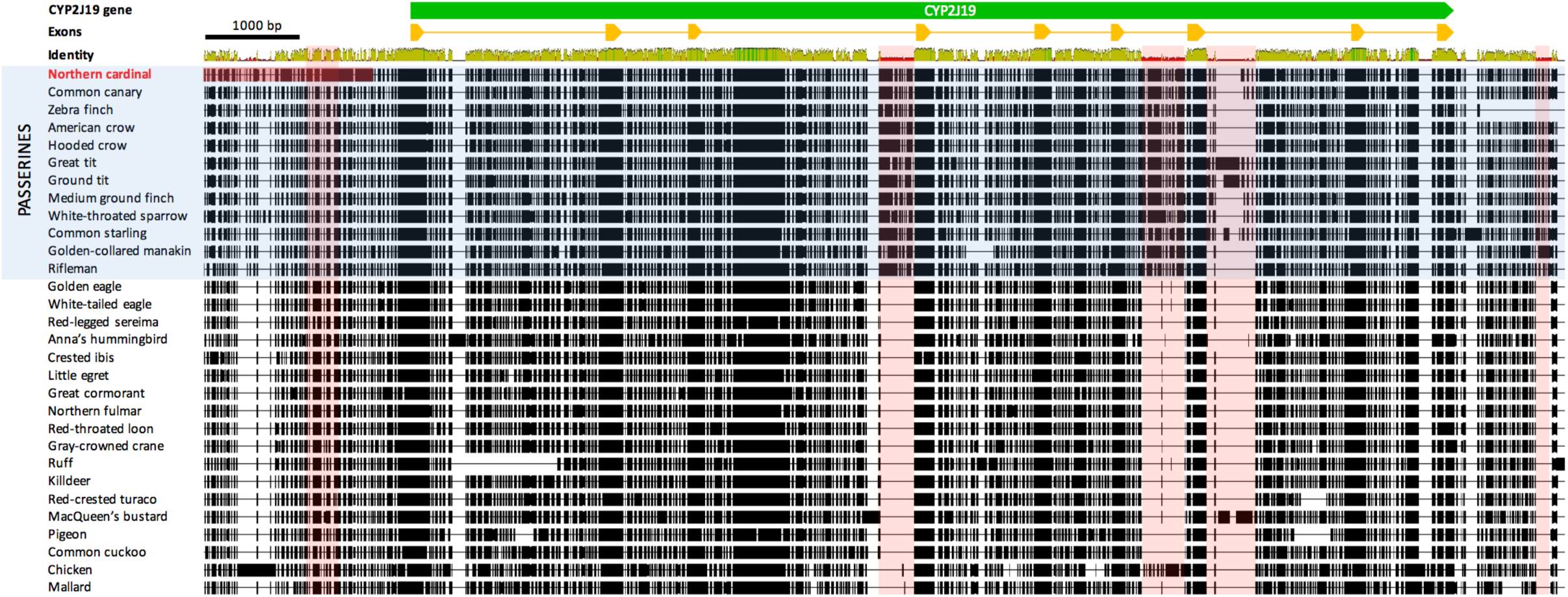
Multiple sequence alignment of the CYP2J19 gene in birds, with red highlighted areas showing regions present in passerines but not other birds. Exons of the CYP2J19 gene are depicted as yellow triangles. The dark red highlighted region upstream of the CYP2J19 gene in the northern cardinal is different from other passerines (see figure 5).

**Figure 4.**
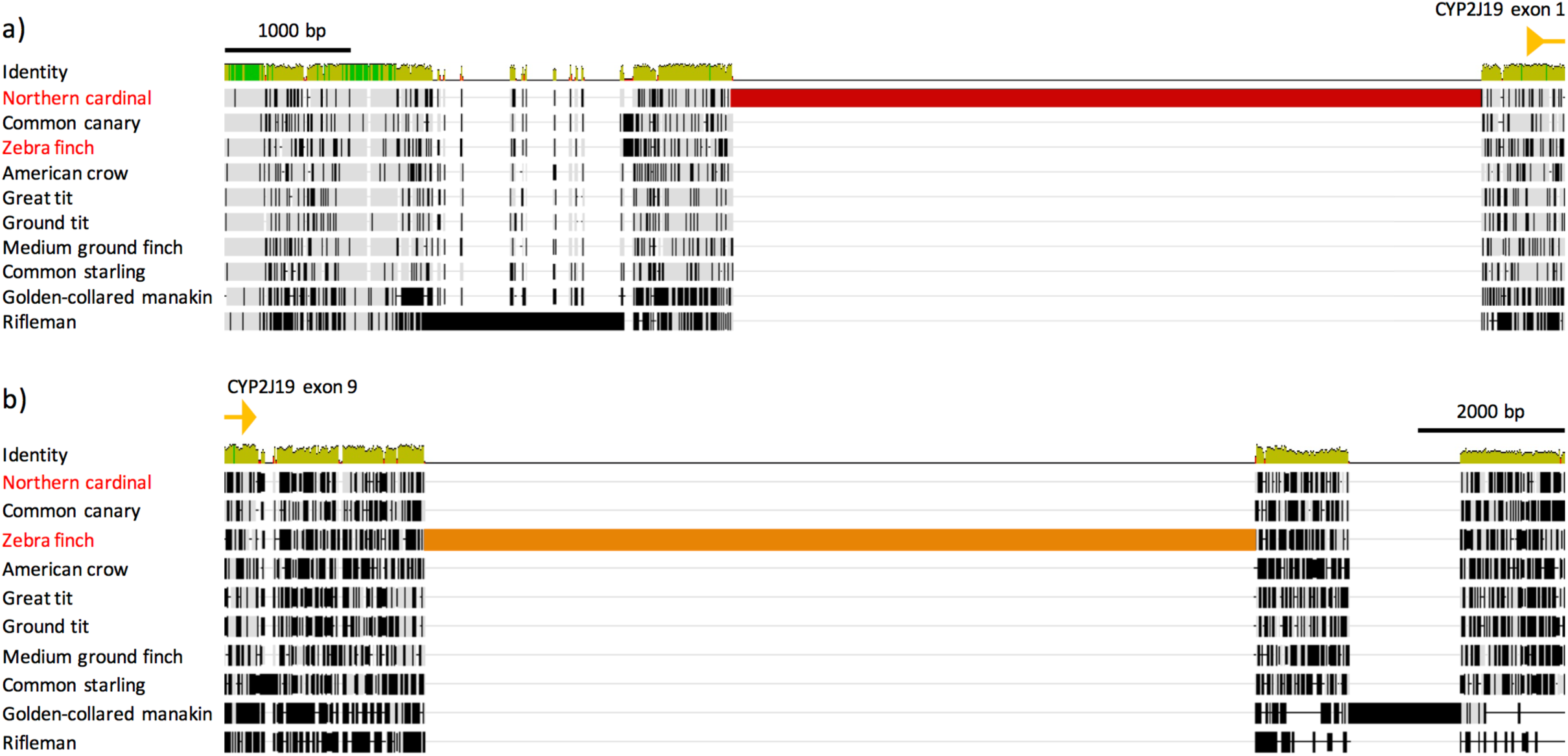
Multiple sequence alignment of (a) upstream and (b) downstream regions of the CYP2J19 gene in passerines. A large insertion upstream of CYP2J19 unique to the northern cardinal is highlighted in red. The insertion of a large region downstream of CYP2J19 highlighted in orange in the zebra finch indicates the CYP2J19B gene in the CYP2J2-like cluster in this species. The names of species possessing red carotenoid coloration (i.e. northern cardinal and zebra finch) are in red. Exons of the CYP2J19 gene are depicted as yellow triangles.

We discovered a total of 25 motifs clustered in a ∼1.7 kb region in the unique insertion sequence of *C. cardinalis* (Fig. 5a). Fourteen TFs are predicted for the 5 identified motif types (Fig. S1, Table 3). No significant GO term appears to be associated with motifs 1–3 and 5, whereas significant GO terms associated with motif 3 predict molecular functions of GTP binding and hexose transmembrane transporter activity. Three predicted TFs (i.e. Sp1, SREBF, and RREB1; Table 3) are associated with androgen regulation of gene expression.

**Table 3.**
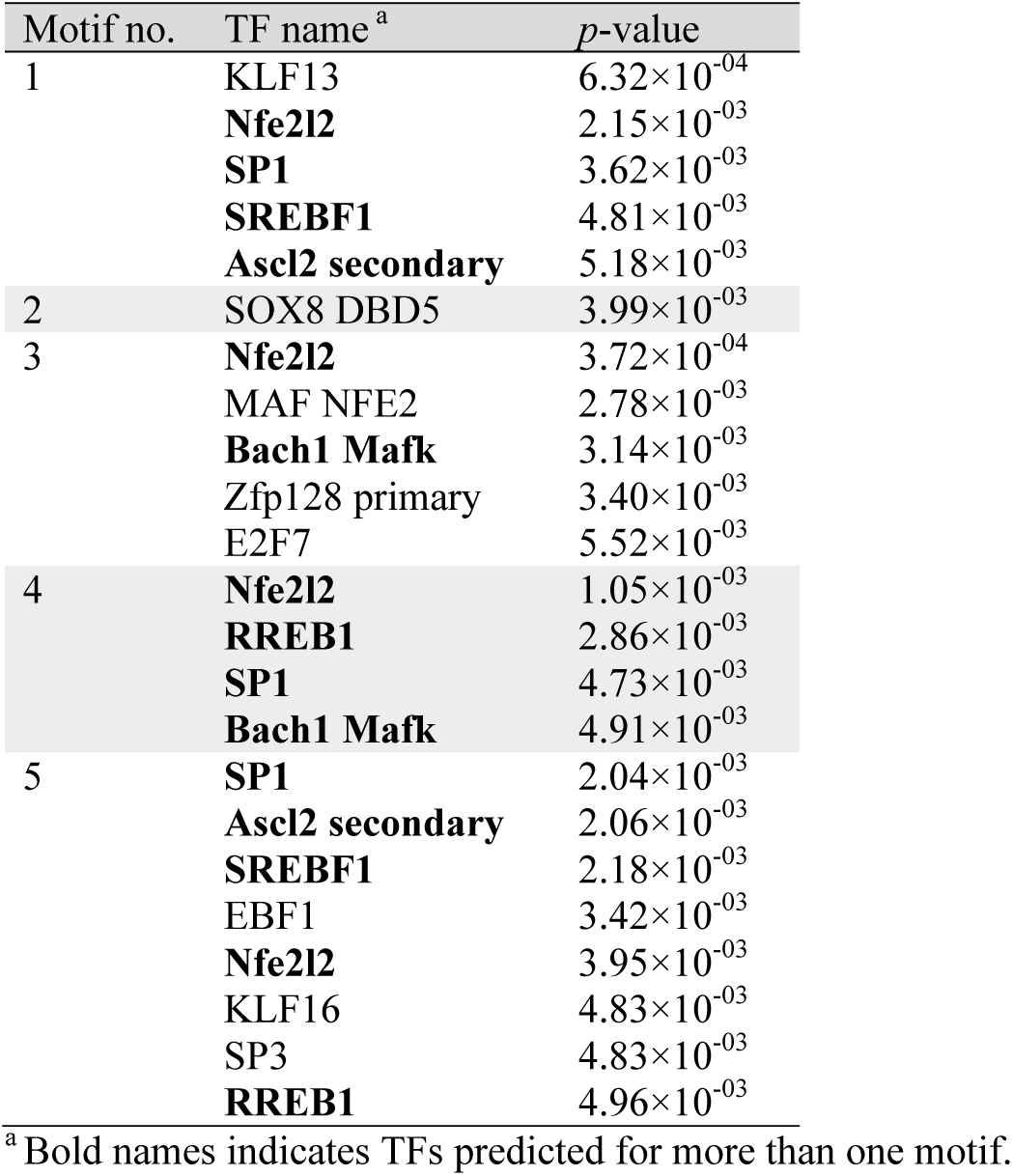
Transcription factors (TFs) predicted for the 5 motifs.

**Figure 5.**
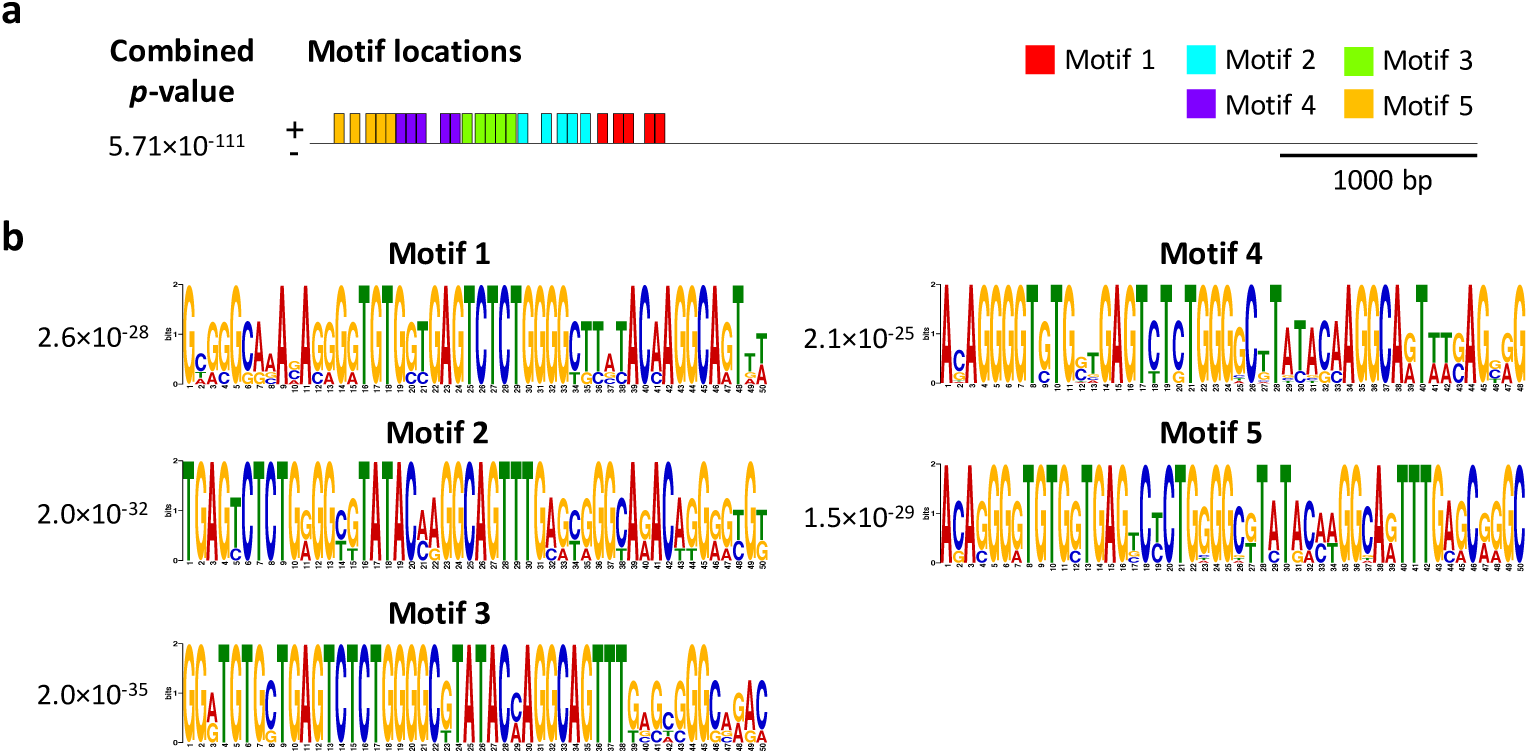
Identification of potential DNA motifs in the unique 5920 bp insertion upstream of the CYP2J19 gene in *C. cardinalis*. (a) Locations of 25 motifs identified and their distribution in the insertion sequence. Sites on the positive (+) strand are shown above the line. Scale is shown below the sequence. (b) Logos of five motifs with statistically significant e-values provided next to the logos.

*Cis*-regulatory elements play a crucial role in the development of sexually dimorphic traits (Williams and Carroll 2009). In birds, sexual dimorphism is strongly affected by steroid sex hormones (Kimball 2006). Sex hormones such as androgen may indirectly influence the expression of androgen-regulated genes, through binding to transcription factors that interact with regulatory elements such as enhancer and cause sex-biased gene expression, which leads to sex-specific phenotypes (Coyne et al. 2008; Mank 2009). Whereas male plumage appears to be testosterone-dependent in passerines (Kimball 2006), the molecular mechanisms controlling sex-biased gene expression and development of sexually dimorphic traits in birds is largely unknown (Kraaijeveld 2019; Gazda et al. in press). In this study, we identified potential TFs that may regulate the expression of the red plumage color gene CYP2J19. Many CYP450s are differentially expressed between the sexes (Rinn and Snyder 2005). Some of our TFs predicted to interact with the insertion unique to *C. cardinalis* are androgen-regulated, including Sp1, which is involved in the androgen activation of the vas deferens protein promoter in mice (Darne et al. 1997) and sterol regulatory element-binding factor (SREBF, aka SREBP), which is involved in androgen-regulated activation mechanisms of target genes (Heemers et al. 2006). Another predicted TF, the zinc finger protein Ras-responsive element binding protein (RREB1), is a co-regulator of androgen receptor (AR) and plays a role in androgenic signaling by affecting AR-dependent transcription (Mukhopadhyay et al. 2007). The interaction between androgens, the implicated TFs and their corresponding binding motifs may underlie the development of sexually dimorphic red plumage color in *C. cardinalis*. In addition, the TF nuclear factor erythroid-derived 2-like 2 (Nfe2l2, also known as nuclear factor erythroid 2-related factor 2, Nrf2) is predicted to bind to all but motif 2 (Fig. 5). The activity of Nrf2 is influenced by changes in oxidative stress, and it regulates the expression of numerous antioxidant proteins (Schultz et al. 2014). Because the production of red ketacarotenoids via CYP2J19 is sensitive to oxidative state (Lopes et al. 2016), this oxidation-dependent TF suggests a possible link between production of red pigments and the oxidative stress faced by the organism, which may reflect an association between red carotenoid coloration and individual quality (Hill and Johnson 2012).

The above non-exonic regions in and outside of the CYP2J19 gene are candidates of regulatory elements responsible for modulating red carotenoid-based coloration in passerines and *C. cardinalis*. Noncoding regulatory regions may be subject to less pleiotropic constraint than protein-coding genes (Carroll et al. 2008), and many CNEEs act as enhancers with regulatory functions (Seki et al. 2017; Sackton et al. 2019). In particular, the large non-exonic region rich with TF-binding motifs upstream of the *C. cardinalis* CYP2J19 gene may play a role in the strong sexual dichromatism and striking bright red male color in this species. The mechanism of plumage color development and sexual dichromatism, and the regulatory role of those genomic regions identified in this study are fruitful areas for future research.

### Demographic historic of *C. cardinalis*

The PSMC analysis (Fig. 6) suggested a demographic history of *C. cardinalis* characterized by a fluctuation in the historic effective population size between 100,000–250,000 around ∼2,000,000 years ago (ya). The population was at ∼160,000 at the beginning of Pleistocene epoch at 2,580,000 ya, then decreased to 110,000 in ∼800,000 ya, and increased to ∼230,000 at the beginning of the Last Glacial Period around 115,000 ya. The population started to declined again after the beginning of the Last Glacial Period (LGP). The decrease in effective population size observed at the beginning of Pleistocene might be due to the divergence of *C. cardinalis* into different subspecies (Provost et al. 2018). The population decline after the start of the LGP is consistent with the ecological niche model that indicates a dramatic range reduction for *C. cardinalis* during the LGP (Smith et al. 2011).

**Figure 6.**
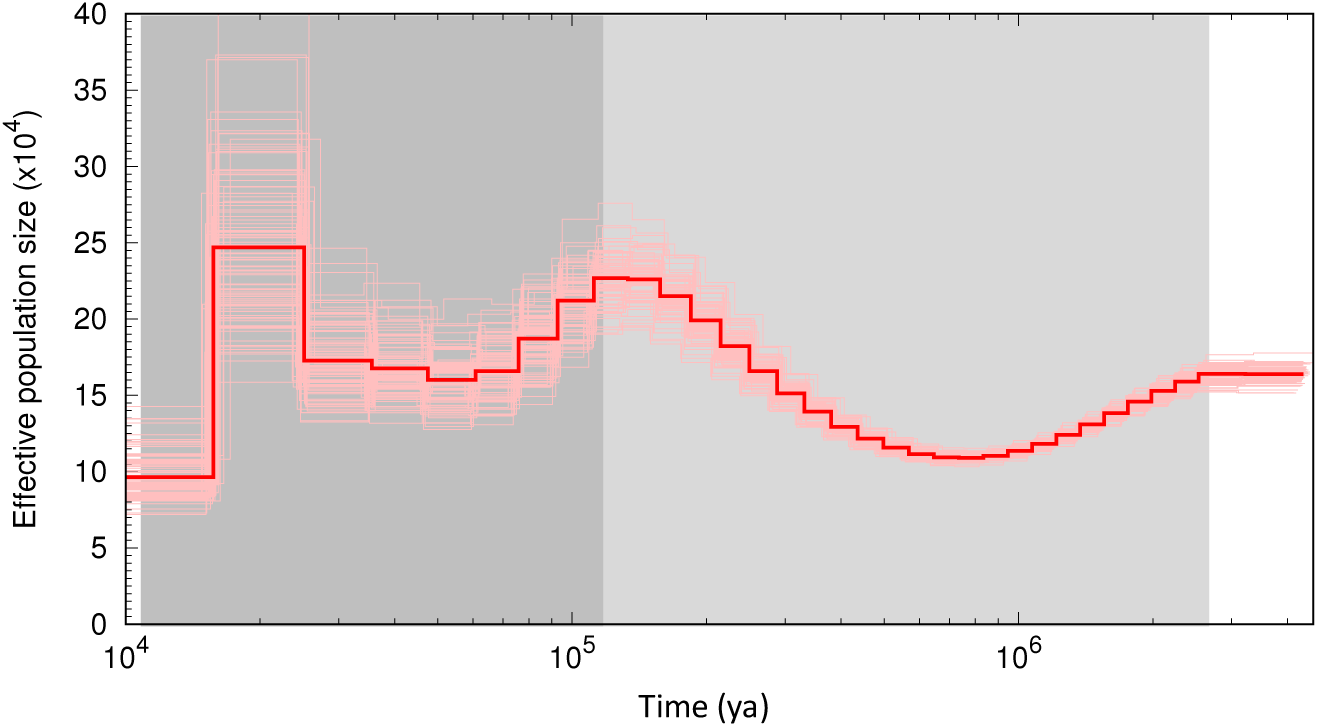
Demographic history of the northern cardinal inferred using PSMC. Bold red line is the median effective population size estimate, whereas thin lines are 100 individual bootstrap replicates. The light and dark grey areas indicate the Pleistocene epoch and Last Glacial Period, respectively.

## Conclusion

We used small-fragment and mate-pair libraries Illumina sequencing data to generate a draft genome assembly of the northern cardinal, *C. cardinalis*. Comparative analyses revealed conserved non-exonic regions unique to the CYP2J19 gene in passerines and in *C. cardinalis*, which may play a role in the regulation of red carotenoid-based plumage coloration. The motifs discovered in the lineage-specific upstream region of CYP2J19 in *C. cardinalis* suggest potential *cis*-regulatory mechanisms underlying sexual dichromatism. The assembled *C. cardinalis* genome is therefore useful for studying genotype–phenotype associations and sexual dichromatism in this species. As one of the first genome sequenced in Cardinalidae, and the highest in quality, it will also be an important resource for the comparative study of plumage color evolution in passerines and birds in general.

## Acknowledgments

We would like to thank Jonathan Schmitt and Katherine Eldridge for collecting and preparing the specimen and tissues. We thank Clarence Stewart for sharing the northern cardinal photo. This research was supported by research funds through Harvard University. The computing resources for this study were provided by the Odyssey cluster supported by the FAS Division of Science, Research Computing Group at Harvard University.

## Supplementary Information

**Figure S1.**
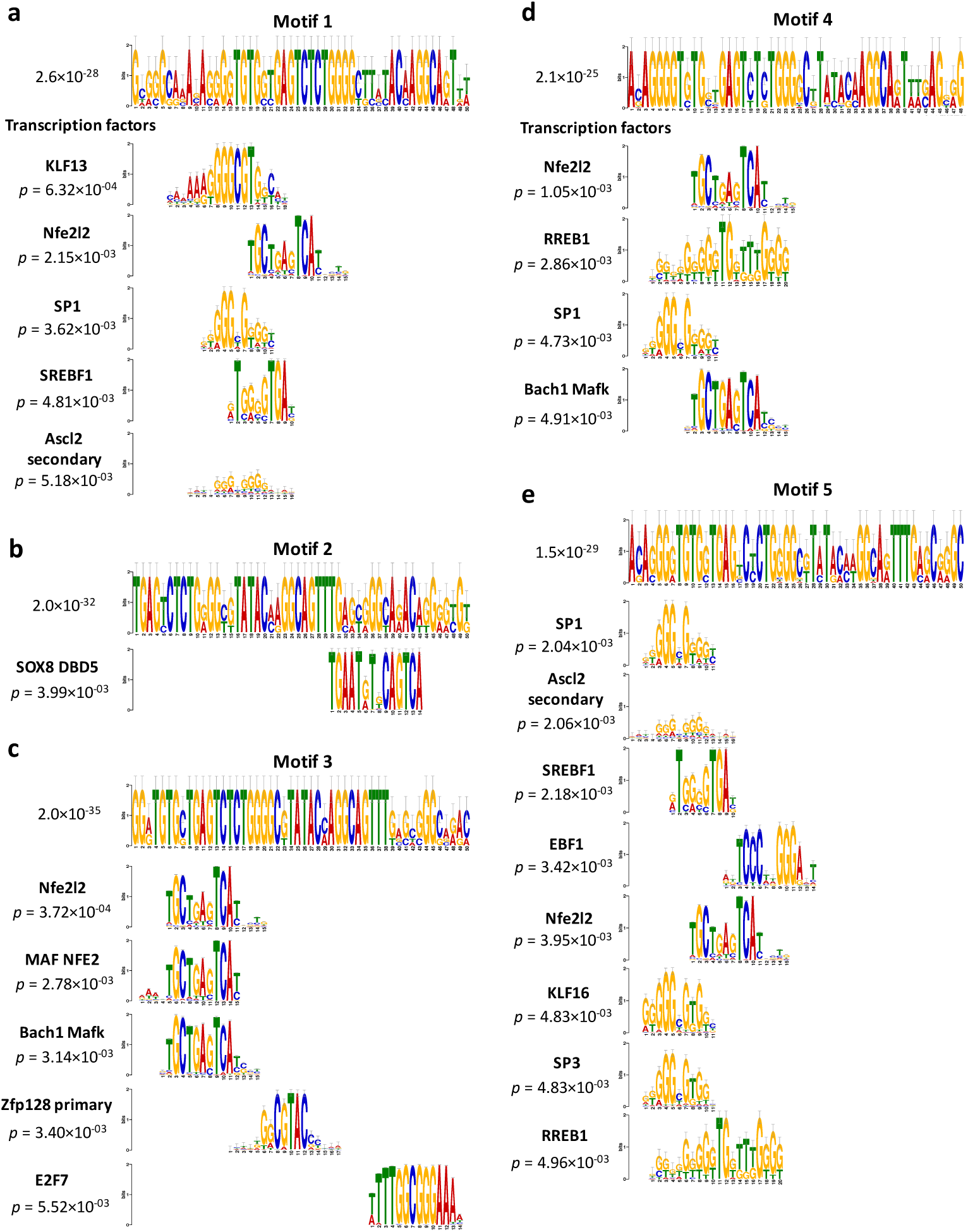
Identification of potential binding transcription factors in the unique 5920 bp insertion upstream of the CYP2J19 gene in *C. cardinalis*. (a–e) Logos of five motifs with statistically significant e-values provided next to the logos. Transcription factors predicted for the 5 motifs are shown below the query motifs.

